# Rocking Without a Tune: *Ex vivo* and *in vivo* Responses to Sound in the Basilar Papilla of the Tokay Gecko

**DOI:** 10.64898/2026.07.16.738899

**Authors:** Brian L. Frost, Yuriria Vázquez, Kazuhiro Horii, Brian A. Fabella, A. J. Hudspeth

## Abstract

Like mammals, some reptiles possess sensitive and frequency-selective hearing at mid- to high-kilohertz frequencies. Both groups employ a place-frequency map within the inner ear to encode sound stimuli, but the mechanisms by which they achieve tuned nerve activation along the cochlea are believed to be distinct. To investigate the mechanical origins of auditory sensation in the reptiles, we measured sound-evoked displacement responses in the hearing organ of the tokay gecko (*Gekko gecko*), known as the basilar papilla. Using optical coherence tomography, we were able to resolve sub-nanometer-scale vibrations throughout the organ both *in vivo* and *ex vivo*. We saw that tuning was not present in the tissue-scale mechanics at any position within the organ; it instead exhibits an untuned rotational motion. We developed a mathematical model which predicted that this motion induces hair-bundle-stimulating fluid displacement, and is due to an anatomical asymmetry seen in many reptile species. These results are consistent with the theory that hair-bundle-level mechanics are primarily responsible for tuning in the tokay gecko cochlea, and we argue that our proposed mechanisms generalize to many other species across the *Reptilia* class.

**SIGNIFICANCE:** We present the most comprehensive picture of tissue-scale mechanics in a reptile hearing organ to-date: sound-evoked displacement responses within the basilar papilla of the tokay gecko (*Gekko gecko*) both ex *vivo* and in *vivo*, across the animal’s auditory frequency range, and along all three spatial dimensions. We observed that place-frequency tuning is not present in the mechanics of the tissue as it is in mammals. Instead, this organ relies on an untuned rocking mechanism to stimulate hair bundles. A mathematical model shows that this may be at play in other reptile species, indicating a unifying mechanism of hearing in the class *Reptilia* in which hair bundle mechanics play the leading role in frequency tuning.

## INTRODUCTION

Hearing is used in predator detection and communication across a large variety of species. Many vertebrates contain a specialized organ within the inner ear known as the *cochlea*, which is responsible for the transduction of sound stimuli into representative nerve responses. This is accomplished by sensory *hair cells*, which are mechanoreceptors topped with *hair bundles* whose stereocilia are arranged in a staircase pattern. Displacement of the bundle toward the tallest row of stereocilia depolarizes the cell, and triggers neurotransmitter release to its afferent contacts (1, 2). Different hair cells in the cochlea respond maximally at different frequencies, and thus the cochlea is capable of breaking down sound stimuli into their component frequencies.

Hair bundles themselves can exhibit resonance, vibrating maximally at a specific frequency determined by length and stiffness. However, this intrinsic tuning alone is not thought to be sufficient for the sensitive, high-frequency hearing characteristic of mammals (3). Similarly, hair cells are electrically tuned (4–10). Although electrical tuning at 4-5 kHz has been hypothesized in some birds (11) and seen in the barn owl (*Tyto alba*) (12), it is generally considered a low-frequency phenomenon; it has only been measured at frequencies below 1-2 kHz in turtles and geckos (9, 13). To achieve high-frequency hearing, hair cells are “helped” by the mechanics of surrounding structures. Within the mammalian cochlea, hair cells are distributed along the cochlear partition, a set of cells and acellular membranes whose size and stiffness vary along the cochlea’s length (14). This impedance gradient means that tissue-scale mechanical responses are tuned to different frequencies at different positions along the cochlea, a phenomenon called *tonotopy*, in which frequency (*tono*) is encoded in space (*topy*) (15–18). Sound stimulation induces a traveling wave along the tissue of the cochlear partition that peaks at a position specific to the stimulus frequency. This tissue-scale mechanical tonotopy displaces hair bundles in a frequency-specific manner, which in turn gives rise to a tonotopy in the nerve response that very closely matches the mechanical response (17). Although hair cell-scale mechanisms may amplify and sharpen this tuning (19–22), the underlying tissue-scale tonotopy is present even in the absence of healthy hair cells (15, 23).

It is known that tonotopy in the nerve response is also present in various reptile cochleae^1^ (26–31) – the reptilian *basilar papilla* (BP, the homologue of the cochlear partition) similarly contains hair cells along its length that produce maximal nerve fiber responses at specific frequencies – but the mechanical tissue-scale tonotopy seen in mammals has not been found in reptiles (26, 31, 32). It is thus hypothesized that tuning arises primarily at the level of the hair bundle (28, 31, 33–39). This hypothesis has been strengthened by recent results in the crested gecko (*Correlophus ciliatus*) basilar papilla, which show that mechanical hair bundle resonance varies along the organ’s length (13, 37). Still, it is an open question as to if or how this hair bundle displacement may be activated by mechanics at the tissue scale; a question complicated by the vast inter-species diversity of reptile BP anatomy (29, 30, 40).

In the present work, we investigated the mechanisms behind tuning and hair bundle stimulation in reptilian ears by observing sound-evoked responses in the BP of the tokay gecko (*Gekko gecko*). These vocal reptiles can hear at relatively high frequencies (up to 6 kHz) and have well-characterized auditory nerve responses (27, 30, 40, 41). While no single cochlea is representative of those of all reptiles, the tokay gecko is a valuable model because it possesses sensitivity over a relatively large frequency range, as well as anatomical diversity within its own hearing organ.

The cochlea of this animal is stimulated by a single middle ear bone (the columella) and is lineal rather than coiled (Fig. 1A). Opposite to mammals, the hair cells at the base of the cochlea are responsible for sensing low-frequency sounds while those at the apex respond to high-frequency sounds (27) (Fig. 1B). The anatomy of the BP varies considerably along its length of only *~1*.7 mm (40). At the base, it is thinner, having a lower density of hair cells whose bundles are covered by a continuous tectorial curtain attached to the limbic cartilage. All basal hair bundles are oriented in the same direction, with their tallest stereocilia facing the neural side. At the apex, there is a higher density of hair cells loaded by distinct tectorial structures: hair bundles on the neural (medial) side are still covered by the tectorial curtain, but abneural (lateral) bundles are connected to discrete sallets that do not attach to any cartilage (Fig. 1A). Apical hair bundles are bidirectional, with some being oriented neurally and others abneurally.

**Figure 1.**
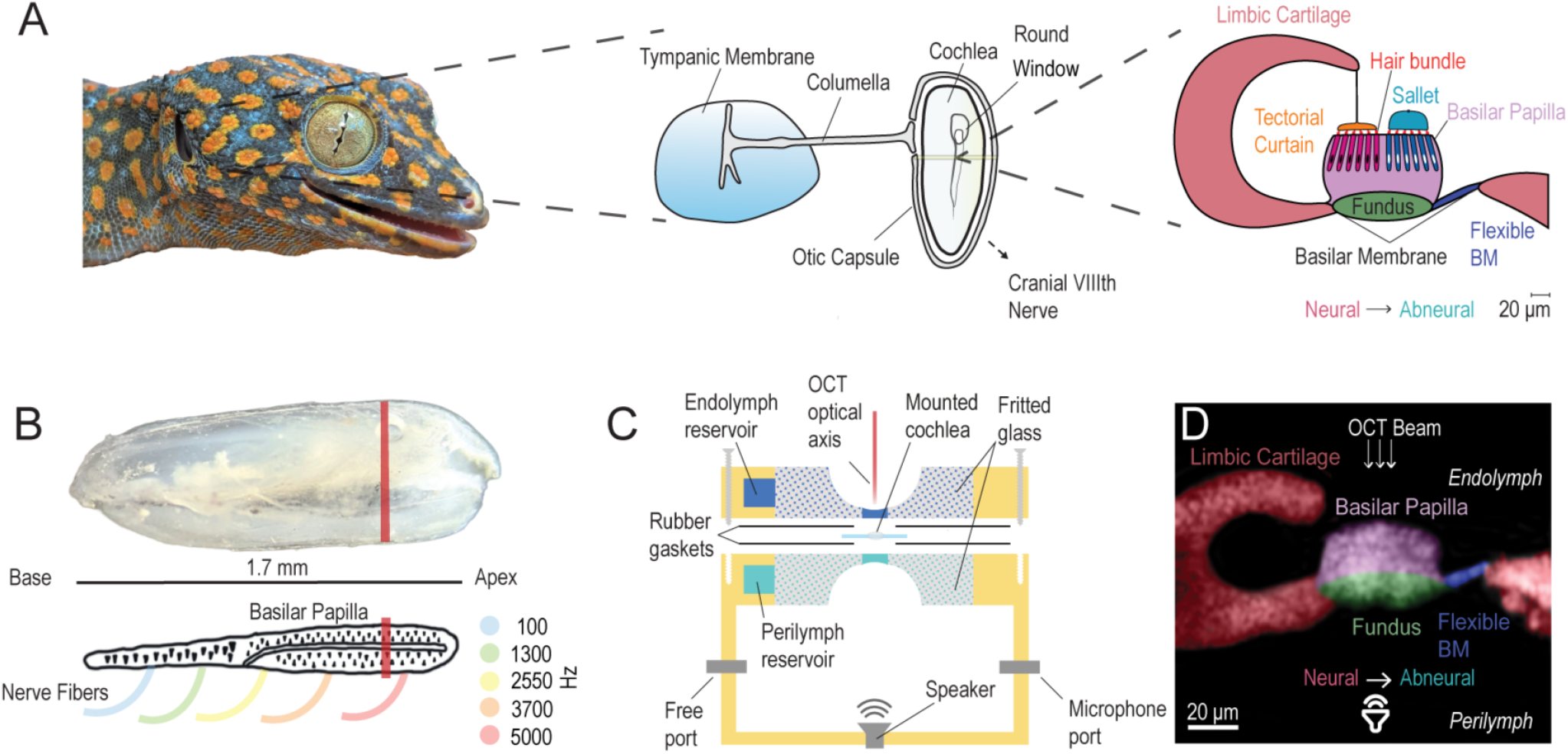
Overview of the tokay gecko ear anatomy and *ex vivo* experimental preparation. (A) Anatomy of the tokay gecko ear. Sound enters the external ear opening (left panel, lateral aspect of the head), mechanically stimulating the tympanic membrane and columella bone of the middle ear, which in turn stimulates the cochlea (middle panel). The cochlea is lined with the sensory epithelium: the Basilar Papilla (BP, an apical cross-section of which is shown in the right panel). Hair cells and supporting cells rest on an acellular structure known as the basilar membrane (BM), comprising a thick portion on the neural side called the fundus and a thinner portion on the abneural side known as the flexible BM. Neural hair bundles connect to the tectorial curtain – a membrane attached to the limbic cartilage – whereas abneural hair bundles are loaded by discrete tectorial structures called sallets. In the basal third of the cochlea, on the other hand, the curtain covers all cells and sallets are not present (40). (B) An excised cochlea from a tokay gecko with the columella and Reissner’s membrane removed, and a corresponding schematic of the BP below. The direction from base-to-apex – the tonotopic axis – is marked. To illustrate tonotopy, curves representing nerve fibers are drawn at five positions along the length of the schematic. Each fiber is colored in correspondence with its best frequency, i.e. the frequency at which nerve responses are maximal at that position (according to Manley (27); the fibers related to the base are tuned to lower frequencies and the fibers related to the apex are tuned to higher frequencies, opposite to the mammal (15, 66)). The orange lines across the cochlea and schematic indicate the approximate position and orientation of the slices seen in panels A and D. (C) Illustration of the two-chamber preparation in which the cochlea is mounted (adapted from Alonso, Gianoli *et al*. (19) with consent of the authors). The corresponding fluid concentrations of endolymph and perilymph were maintained within the chamber while the organ was acoustically stimulated and displacement responses were measured. The beam of the OCT device points into the cochlea on the endolymph side for imaging and displacement measurements. Sound stimulation is provided on the perilymph side, and a microphone is situated within the chamber to observe precise stimulus pressures. (D) A labeled OCT image of a cross-section of BP, comparable in orientation to the illustration in the right of panel A. As the device produces unlabeled images with a resolution of *~* 4*μm*, landmarks like the limbic cartilage, must be used to infer the anatomical structures being observed. Tectorial structures were not visible in our preparation, but may still be present (see “Open questions and limitations of the study” subsection).

The cochlear fluid spaces are separated by an acellular structure known as the *basilar membrane* (BM) which exhibits an asymmetric geometry – there is a thicker *fundus* on which the hair cells and supporting cells of the BP sit at the medial end, and a thinner portion lateral to the cells that we call the flexible BM (Fig. 1A). This BM asymmetry is present not only in other lepidosauromorphs, but also in testudines and crocodilians, which are known to have quite advanced hearing (30, 40).

Our study offers a view of the sound-evoked mechanics of the tokay gecko cochlea along its three dimensions both *ex vivo* and *in vivo* by using optical coherence tomography (OCT), a technique capable of at-depth imaging and vibrometry at sub-nanometer scales (42–44). In *N* = 29 BPs *ex vivo* and *N* = 4 BPs *in vivo*, we found no position at which responses were tuned to a specific frequency in the animal’s hearing range. Instead, the organ rotates about its tonotopic axis, perpendicular to the hair bundles *–* it *rocks* side-to-side within the plane of Figure 1A or D, with abneural and neural sides moving half of a cycle out of phase with one another *–* in an untuned manner. This untuned rocking is present across the organ’s tonotopic axis, i.e. at locations responsible for the sensation of both higher and lower frequencies. Guided by the results of a mathematical model, we hypothesize that this untuned rocking is caused by the aforementioned BM asymmetry, and results in fluid being pushed in the hair-bundle-stimulating direction, activating a tuned mechanism therein.

An untuned rotational mode of motion, as we have seen in this organ, is known to be present in the BPs of other lizard species (32, 36), and has been suggested by computational models to be present in the BP of the tokay gecko (34, 39). Our results both validate and extend this long-standing theory, as 1) we are able to measure the displacement across the entire span of the BP in three dimensions using OCT, 2) we are able to achieve *in vivo* recordings, and 3) the intra-organ diversity of the BP in this species offers insight into other reptilian hearing organs.

In particular, the robustness of untuned rocking across both low- and high-frequency regions – and thereby salletal and non-salletal regions – in this organ, as well as the presence of the same BM asymmetry in many reptile BPs (30, 40), indicate that this mechanism could be employed in a large variety of related species. For example, the results are qualitatively similar to previously measured responses at the surface of the alligator lizard BP (26, 32), a species with a shorter cochlea lacking either of the tectorial structures present in the tokay gecko, instead having free-standing hair bundles.

Our results are consistent with the theory that tuning in the reptile ear is primarily the result of hair bundle resonance, evincing that hair bundle mechanics can play a significant role in the function of a hearing organ sensitive to high frequencies.

## RESULTS

Tokay gecko cochleae were isolated and placed in a modified version of a two-compartment chamber described previously (19) (Fig. 1C, see Methods section for more details). Using OCT, we obtained at-depth images of the cochlea, such as the one shown in Figure 1D, whose orientation is similar to the illustration in Figure 1A. Across all OCT images, the C-shaped limbic cartilage (on the upper-left in these images) served as a primary landmark for orientation. Along with imaging, OCT is also capable of measuring vibrations at sub-nanometer scales at sufficiently bright pixels within the image (43); we used this to record displacement in response to sound stimuli across the animal’s range of hearing (500-6000 Hz (27, 40)) in *N* = 29 organs.

We analyzed these signals in two ways: by examining responses at individual positions within the organ as a function of the stimulus frequency, and by examining responses across an entire cross-section of the organ at a single stimulus frequency. The first approach allowed us to determine whether the basilar papilla exhibits tonotopy at the tissue scale, as described in the following subsection. The second revealed a rocking motion of the organ, presented thereafter.

Similar data were then acquired *in vivo* in *N* = 4 preparations, showing consistency with our results from the excised preparation.

### Lack of tissue-scale mechanical tonotopy in the tokay gecko cochlea

We observed displacement responses across *N* = 29 BPs and found that no position within the organ was tuned to a specific frequency. Instead, responses were mildly low-pass across the known range of tokay gecko hearing.

Figure 2 shows frequency responses from a single tokay gecko cochlea at several positions to multitone stimuli (45, 46) within the animal’s hearing range (13 tones, each at 80 dB SPL^2^, between 531 Hz and 6078 Hz; see the Methods section for more details). At each of three positions along the cochlea’s length, we present the sensitivity (displacement normalized by stimulus pressure) as a function of frequency at three anatomical locations: the BM, the abneural hair cell region (lateral, or “right” in the image) and the neural hair cell region (medial, or “left” in the image).

**Figure 2.**
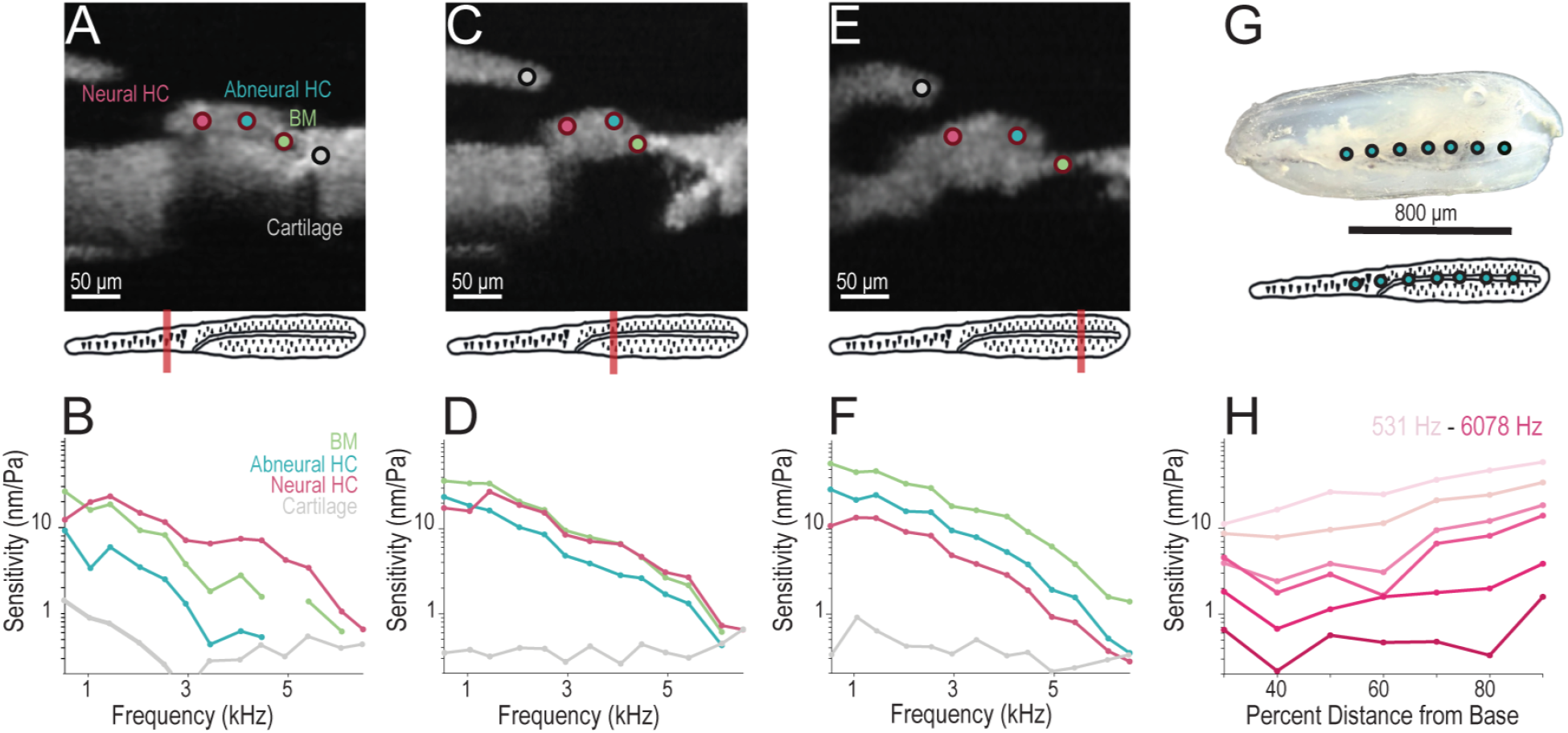
Evidence for the lack of tissue-scale mechanical tonotopy in the tokay gecko cochlea. (A) OCT image of the basilar papilla of the tokay gecko cochlea at a basal location (TK87, 30% from the base). Colored dots indicate locations at which sensitivity responses are shown in the next panel – one in the neural (more medial) hair cell (HC) region, one in the abneural (more lateral) HC region, one at the basilar membrane (BM), and one in the cartilage that serves as a control. The orange line in the schematic below indicates the approximate position and orientation of the scan within the basilar papilla. (B) Sensitivity (displacement magnitude normalized by stimulus pressure) as a function of stimulus frequency in response to an 80 dB SPL zwuis stimulus (45, 46). Each curve corresponds to the position in panel A marked with a dot of the same color. Note that sensitivities are low-pass rather than being tuned to a specific frequency. Displacements of the limbic cartilage, alternatively, show low-magnitude broadband behavior, indicating that bulk preparation movement is not responsible for the measured intra-BP response. (C-D) Same as A-B but at a more central position in the same cochlea (*~5*0% from the base). Notably, despite being at a different location, mechanical responses look similar to those at the other measured cross-sections. (E-F) Same as A-B but at a more apical position in the same cochlea (*~8*0% from the base). (G) Excised cochlea and corresponding schematic, marked with longitudinal locations at which the displacement responses in the following panel were measured. (H) Sensitivity of the BM as a function of longitudinal position at six distinct frequencies in the same cochlea. Measurement locations are marked in panel G. Notably, there is no discernible tuning to a single position for any of the stimulus frequencies.

In the basal cross-section (Fig. 2A-B), motion in all three regions is seen to be mildly low-pass across the range of hearing. The same result is seen at central (Fig. 2C-D) and apical (Fig. 2E-F) cross-sections. No visible structures show tuning to any frequency, in contrast with known neural tonotopy (27). Instead, responses are maximal in the 500-1500 Hz range and characteristically similar regardless of position within the basilar papilla.

Reciprocally, we observed the response to a single frequency within the neural hair cell region at a variety of longitudinal positions along the same cochlea (Fig. 2G-H). In this case, we observed that no frequency was tuned to any given position. Instead, responses appear to be slightly larger at the apex than at the base regardless of stimulus frequency.

Thus, the two elements of tonotopy – frequencies tuning to positions, and positions tuning to frequencies – are not observed in the tissue-scale mechanics of the tokay gecko BP.

As a control, we also measured the sensitivity within the cartilage near the organ. The cartilage moves in a broadband fashion, at a low amplitude relative to intra-BP responses (Fig. 2B, D, and F, gray line). This shows that the low-pass nature of our measured BP responses is not merely the result of our stimulus, or bulk low-pass motion of the entire preparation, but indeed the mechanics of the organ itself.

### Antiphase “rocking” motion in the basilar papilla

We measured the sound-evoked displacement response across entire cross-sections of the basilar papilla, and found a characteristic antiphase pattern: the neural and abneural cells move approximately half of a cycle (180^°^) out of phase with one another. The patterns are represented as color maps in Figure 3A, C, and E, where saturation encodes magnitude and hue encodes phase.

**Figure 3.**
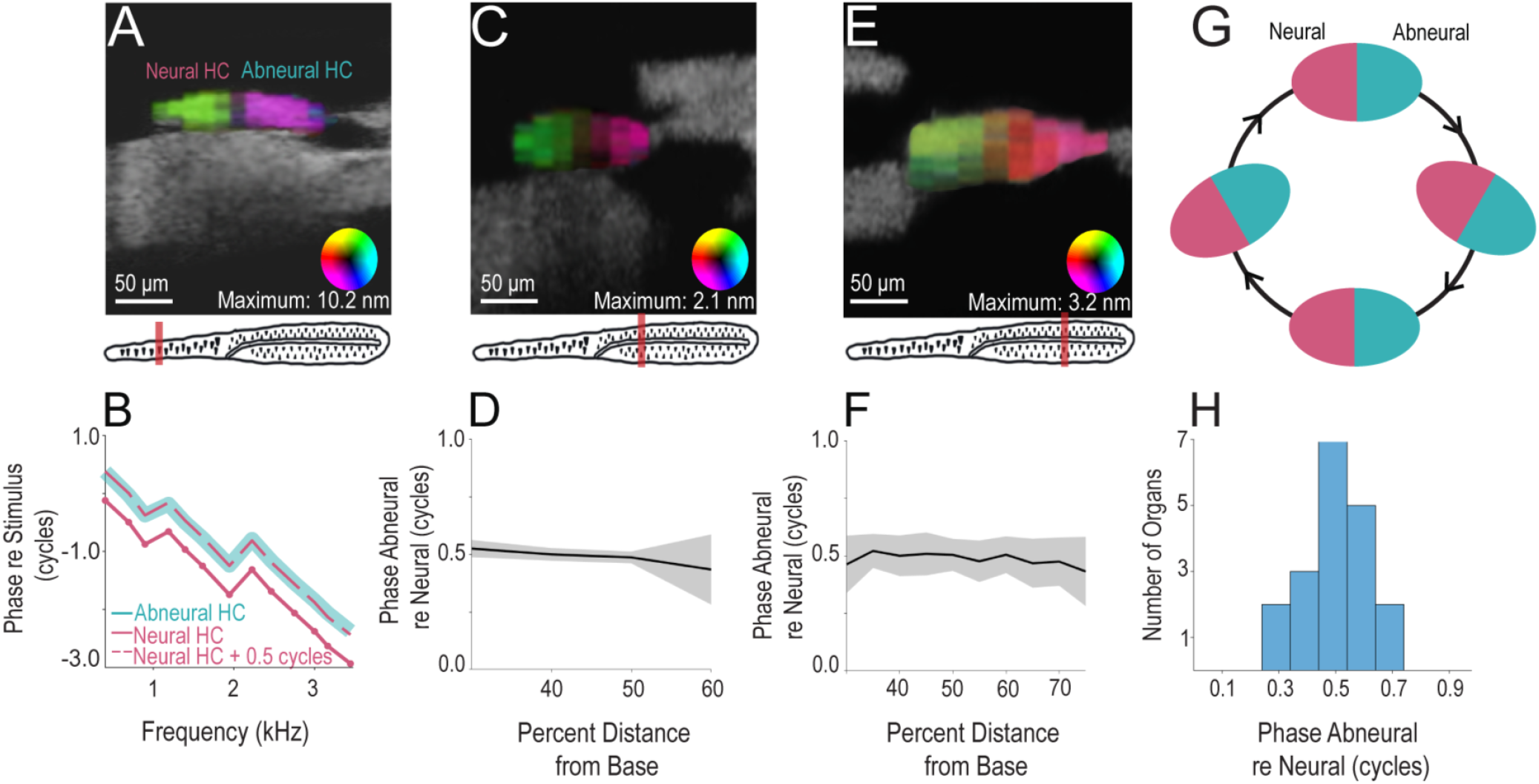
The organ rocks: antiphase motion within the organ at various tonotopic positions and frequencies. (A) OCT image from the basal region of the cochlea (TK 19, *~* 20% from the base) with a heat map encoding displacement (80 dB SPL, 2200 Hz stimulus) overlaid on the BP. In the colored overlay, saturation encodes displacement magnitude (maximum 10.2 nm), while hue encodes phase relative to sound stimulus. The color wheel at the bottom-right of the panel is used to interpret the coloring – for example, the right of the color wheel (teal) represents a response in-phase with the stimulus, while the left of the color wheel (red) represents a response one half-cycle out of phase with the stimulus. (B) Phase of hair cell displacement relative to stimulus pressure as a function of frequency. Solid curves are responses at positions within either the neural or abneural half of the organ. The dashed line represents the neural-side response shifted upwards by 0.5 cycles; it lies nearly on top of the abneural-side response (maximum absolute difference of 0.07 cycles), illustrating that the sides move approximately in antiphase. (C) Similar to A but in a central region of a different cochlea and with a different stimulus frequency (TK 77, *~6*0% from the base, 1042 Hz). (D) Phase response at a position in the abneural region (*ϕ*_*A*_) relative to that of a position in the neural region (*ϕ*_*N*_) at four positions along the length of the cochlea (TK 77, as in panel C). The solid line is the average of this quantity over nine different stimulus frequencies (500 to 5000 Hz, spaced near-evenly); shaded region denotes one standard deviation from the mean. Note that at each position, the two sides move approximately 0.5 cycles apart. (E) Same as A and C but at a more apical position of a different cochlea and with a different stimulus frequency (TK 84, *~8*0% from the base, 3443 Hz). (F) Same as D, but at 11 positions along the length of the cochlea (TK 84, as in panel E). Averaged over ten frequencies between 500 Hz and 6 kHz, spaced near-evenly. (G) Schematic illustrating the organ’s motion in response to a single tone at different points in a cycle if the two sides move precisely in antiphase; the organ rocks side-to-side. Displacements are in reality very small compared to the size of the organ, but are exaggerated here for illustrative purposes. (H) Histogram showing the phase difference of abneural re neural hair cell displacement at a frequency between 1 kHz and 2 kHz in 19 organs. The phase difference is 0.50 ± 0.12 cycles (mean ± standard deviation; N=19).

Observing the response across a basal cross-section at 2200 Hz, we see a striking near-binary nature of the phase response, where the two sides of the organ respond at similar magnitudes but in antiphase (Fig. 3A). This principle extends across the hearing range of the animal (Fig. 3B): the phase responses at one position in the abneural hair cell region and one position in the neural hair cell region were plotted as a function of frequency, as was the neural response shifted by one half of a cycle, and the abneural and shifted neural phase responses lie nearly on top of one another. The two differ by at most 0.07 cycles (25^°^) across presented stimulus frequencies.

In two organs, we were able to measure displacement at both neural and abneural positions at various positions along the tonotopic axis (base-to-apex). We averaged the phase difference between neural and abneural positions across frequency, and found the average to be about half of a cycle regardless of tonotopic position (Fig. 3D, with an example cross-section of this organ shown in panel C) This was repeated along a larger span of a second organ, confirming the same result (Fig. 3 E-F).

We conclude that this antiphase motion is robust in frequency and tonotopic position. More examples are shown in supplemental Figure S1.

The displacement measured by OCT vibrometry is a projection onto the optical axis (48–50), so this means one side of the organ is moving towards the OCT device (“up” in the image) while the other is moving away from it (“down” in the image). Assuming the optic axis is normal to the BM (approximately the case in our preparation), Figure 3G is a schematic of how antiphase motion manifests in the time domain, with one side moving in the opposite direction to the other; the organ appears to rotate side-to-side. In holding with previous literature (32, 36), we refer to this phenomenon as *rocking*.

We were able to assess both neural- and abneural-region phase responses within at least one cross-section of *N* = 19 BPs. We considered the phase difference between one neural and one abneural pixel at a single longitudinal position in each BP in response to the tone between 1 kHz and 2 kHz at which the SNR was highest in that organ (Fig. 3H). The distribution shows that the phase difference is tightly centered around a half-cycle – the phase difference was 0.50 ± 0.12 cycles (mean ± standard deviation; *N* = 19). Thus, rocking motion is consistent across organs.

### Displacement responses *in vivo* are consistent with *ex vivo* results

In *N* = 4 organs, we measured sound-evoked displacement responses of the BP *in vivo* using an OCT device with a higher center wavelength (1300 nm as opposed to 900 nm, offering superior penetration depth at the cost of axial resolution (42)). By making an incision in the lower jaw, we removed muscles and tissue to expose the cochlea’s round window membrane; this membrane was then opened, and the animal was oriented so that the BP in the apical region could be imaged by OCT through the round window (Fig. 4A). Sound was provided by a speaker placed near the outer ear opening ipsilateral to the cochlea from which displacement responses were measured. The schematic in Figure 4A illustrates the preparation; additional experimental and surgical details are presented in the Methods section.

**Figure 4.**
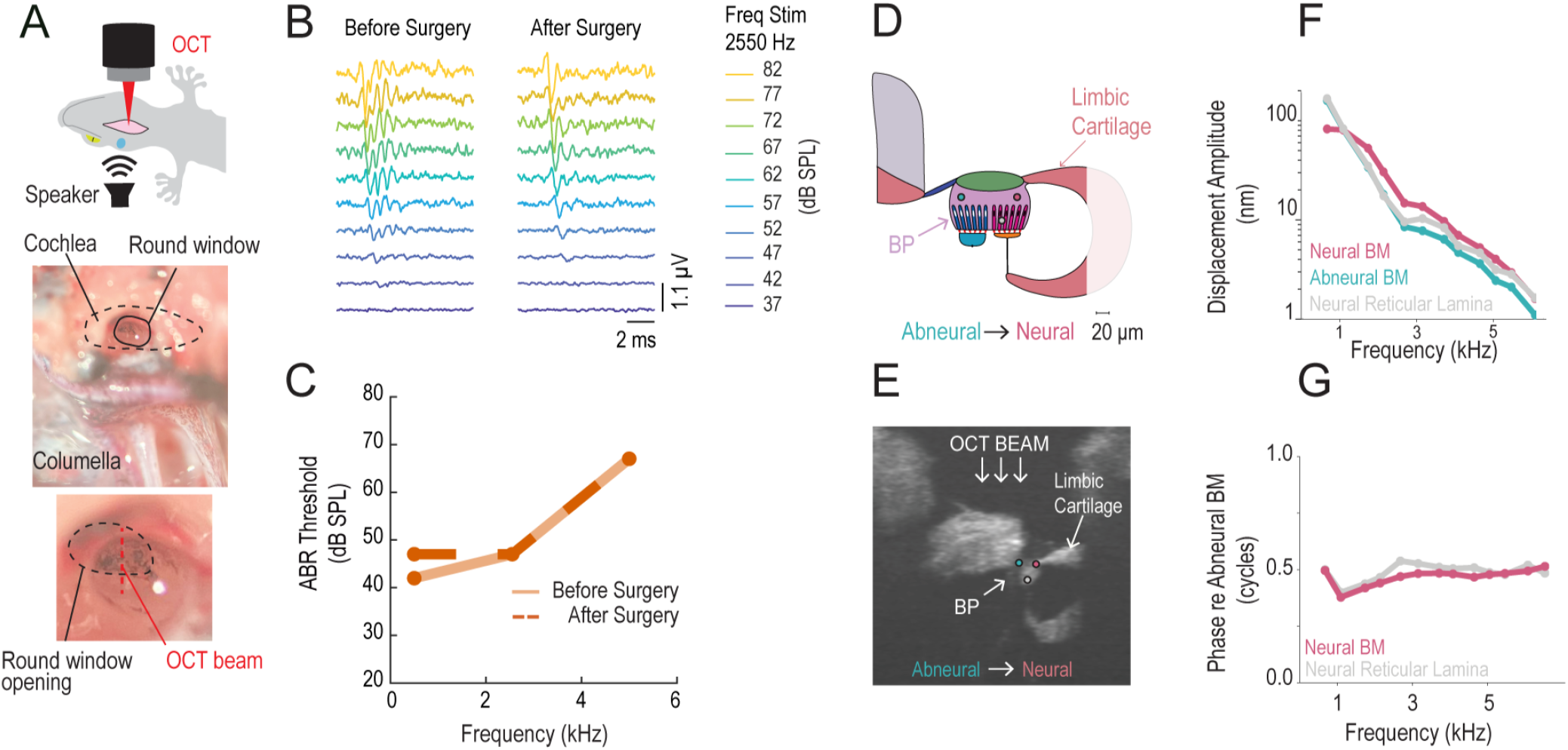
In vivo OCT measurements of the tokay gecko organ. (A) Schematic of the surgical preparation, acoustic stimulation, and OCT measurement in the tokay gecko *in vivo*; the organ is viewed through the opened round window, exposed by an incision in the lower jaw and excision of the muscle and soft tissue surrounding the cochlea. Acoustic stimulation was provided through a speaker placed near the external ear opening on the ipsilateral side of the cochlea from which images and displacement responses were measured. (B) Representative ABR traces to an acoustic stimulus at 2550 Hz for an example gecko (TK 99) before and after the surgical procedure. The ABR threshold (the lowest sound amplitude that triggers a neural response) was 47 dB SPL before and after the surgery. (C) ABR thresholds in the same animal as a function of stimulus frequency (500, 2550, 5000 Hz) before and after surgery, showing no significant degradation. (D) Anatomical drawing of the BP at this orientation, with colored circles indicating approximate positions at which three displacement responses were measured: abneural and neural position on the BM and a neural position on the reticular lamina. (E) OCT image of BP *in vivo* in the same animal as viewed through the opening illustrated in panel A. At this orientation, the BM is nearer to the lens – “upside-down” relative to our *ex vivo* recordings. Colored circles represent positions at which displacement responses are displayed in F and G. (F) Displacement amplitude as a function of the frequency in response to an 80 dB zwuis stimulus at the locations marked in D: neural BM, abneural BM, and neural reticular lamina (the surface of the hair cells where the bundles lie). (G) Phase responses at neural BM and neural reticular lamina positions relative to abneural BM phase as a function of stimulus frequency (corresponding to the magnitude responses in F). These neural positions are seen to move approximately one half-cycle out of phase with the abneural position, consistent with a rocking motion.

We assessed cochlear health by measuring the auditory brainstem response (ABR), a non-invasive diagnostic procedure of cochlear and central auditory system function (51, 52) (for details, see the Methods section). ABRs were recorded before and after surgery to confirm that the procedure did not damage the cochlea. If post-surgical ABRs remained and their thresholds were within 25 dB of the baseline at all three frequencies tested, we measured sound-evoked displacement responses; otherwise, the data were discarded.

ABR responses to 2550 Hz in one gecko are shown in Figure 4B; responses before and after the surgery remained similar. Figure 4C shows the thresholds for the three stimulation frequencies within the gecko’s range of hearing before and after the surgery. All ABRs remained and their thresholds were within 5 dB of baseline at all frequencies tested, indicating that the cells within the organ were intact and the pathways between the cochlea and the brain were functioning.

As in the *ex vivo* preparation, we found no position within any of *N* = 4 BPs *in vivo* at which mechanical tuning was present. Instead, responses were low-pass within the range of hearing. Figure 4E shows an OCT image^3^ of one BP in which we measured from the BM at neural and abneural positions, as well as at the reticular lamina *–* the surface of the BP on which the hair bundles lie *–* in a neural region. The displacement responses at these three positions are presented in Figure 4F; the responses are seen to fall off with increasing frequency even in the apical region where the nerve responds to higher frequency. This strongly resembles the *ex vivo* displacement responses of Figure 2. Examples of this behavior from three other animals *in vivo* are shown in supplemental Figure S2.

In one animal, we were able to compare phase responses at the neural and abneural sides of the BM. We found that the two sides move out of phase by about half of a cycle across frequency (Fig. 4G), consistent with the presence of the rocking motion seen in our *ex vivo* preparations (Fig. 3). While we could only record statistically significant displacement responses on the neural side of the reticular lamina, we see that its phase response is also consistent with the data from the excised preparation. Namely, it is approximately in-phase with neural-side BM motion, and thereby about a half-cycle out of phase with abneural-side BM motion (Fig. 4G).

The constraints of the *in vivo* preparation made measurements across the radial width of the organ and within the cells of the BP more challenging to attain. Still, these preliminary results suggest that our two *ex vivo* findings – the lack of tonotopy and the presence of rocking motion – are consistent with *in vivo* physiology.

### A two-beam model for the basilar membrane explains rocking and hair bundle stimulation

A model was developed to investigate the origin of the antiphasic rocking motion in the gecko cochlea seen in Figure 3, and to explore the consequences of this displacement pattern for hair bundle stimulation.

The BM of the tokay gecko is split into two components: a thick and round fundus and a thinner flexible BM (40) (Figure 5A, a zoom-in of the OCT image in Figure 1D). The fundus is significantly thicker than the flexible BM, so it is expected to be stiffer. We modeled the BM at a single tonotopic position as two beams of different lengths and stiffnesses, with the stiffer medial beam representing the fundus and the more compliant lateral beam representing the flexible BM (Fig. 5B). Using the dynamic beam equation, we simulated the displacement of this composite beam system in response to a sinusoidal pressure stimulus. We then solved for the fluid displacement generated by a membrane moving in such a way at one edge of a fluid-filled compartment. Mathematical details are given in the Methods section and supplemental information.

**Figure 5.**
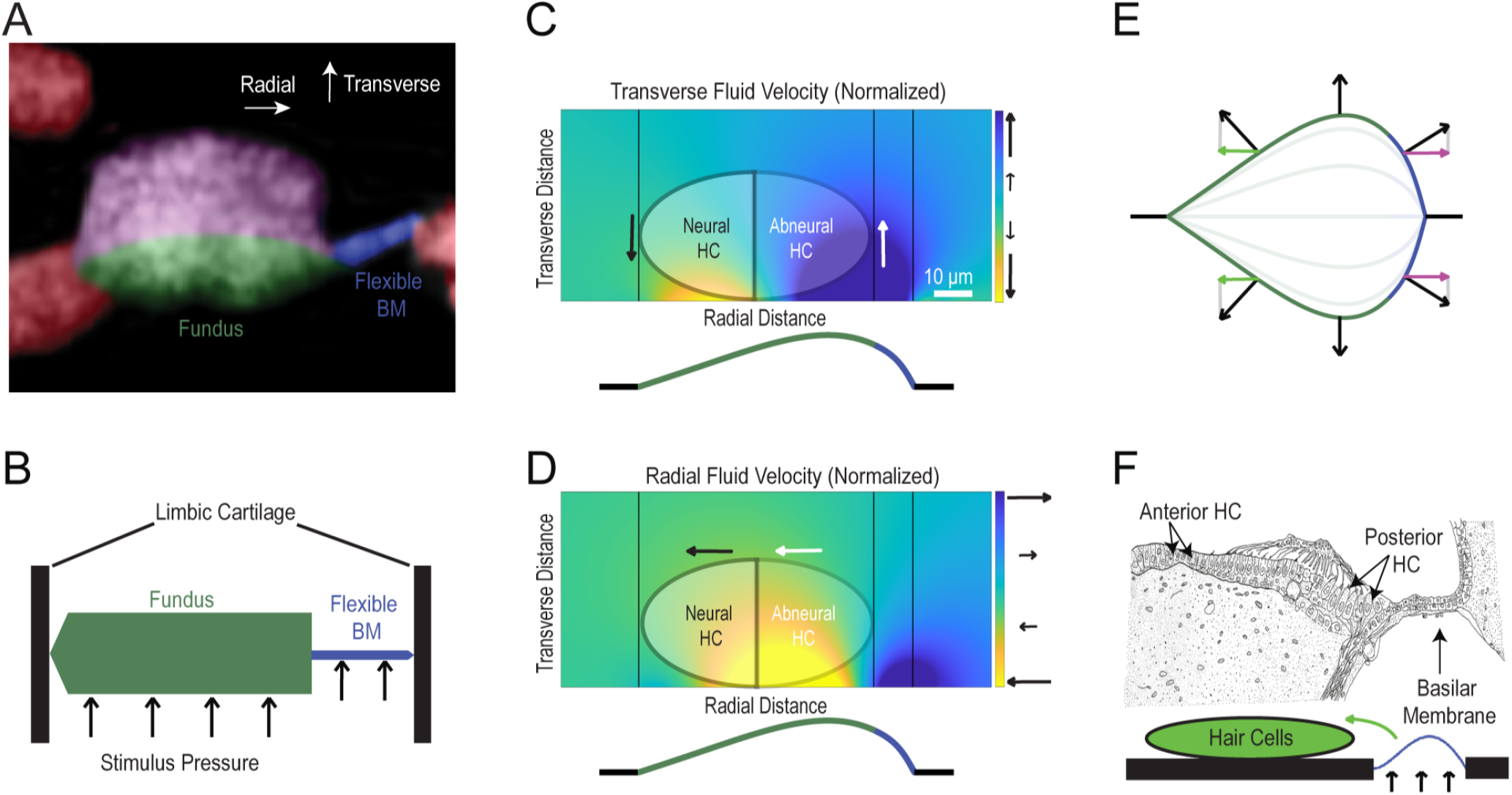
Results and predictions from the two-beam model. (A) An OCT image of the BP from Figure 1D, zoomed in to emphasize the structure of the BM. The anatomical axes – transverse and radial, normal and parallel to the BM, respectively – are noted on the top of the image. (B) A schematic of the two-beam model of the BM inspired by this OCT image. The fundus and flexible BM are modeled as two joined beams, with their non-mutual ends simply supported by the limbic cartilage. The fundus is modeled as being longer, thicker and stiffer than the flexible BM. (C) Modeled transverse (“up-down”) fluid velocity driven by the BM motion computed from the two-beam model, at one point in time in response to a uniform sinusoidal pressure stimulus. Approximate positions of hair cells (HCs) are shown overlain, and arrows are used to represent the predicted directions of motion of the abneural and neural hair cells in this direction. Note that the sides of the organ move opposite one another. Black vertical lines indicate the joints between (left-to-right) cartilage and fundus, fundus and flexible BM, and flexible BM and cartilage. A representation of BM displacement is shown below the image color-coded by region – black for cartilage, green for fundus and blue for flexible BM. (D) Similar to panel C, but showing the radial (“left-right”) component of displacement. In this direction, both neural and abneural HC regions are stimulated in-phase with one another. (E) Modeled displacement response to a sinusoidal pressure, showing the asymmetric motion of the BM. Responses are shown at every third of a cycle, with opaque curves showing displacement at the peak and trough of the cycle. Displacements are in reality very small compared to the size of the organ, but are exaggerated here for illustrative purposes. Normal vectors to the BM are shown to illustrate the direction in which fluid immediately adjacent to the BM might be pushed; the normal points medially on the medial end and laterally on the lateral end. (F) Illustration of the BP of the red-eared slider turtle (*Chrysemys scripta elegans*) adapted from *Wever (1978)* (40). This is an extreme case of the presented model, where the hair cells lie on the cartilage rather than on a stiff fundus. The schematic indicates how having a flexible BM lateral to the hair cells may explain how hair bundles are stimulated in species with such BP architectures.

Figure 5C-D shows the predicted fluid velocity for this model, both in the transverse (up-down, panel C) and radial (left-right, panel D) directions as colored maps. One assumption of our model is that the cells behave mechanically as fluid, and they are thus not explicitly modeled (as in the model of Aranyosi and Freeman (32)). This is informed by: 1) the BP does not contain the complex cytoarchitecture of stiff cells seen in the mammalian organ of Corti that distinguish its motion from that of fluid (14, 30, 40, 53–55), and 2) it does not exhibit somatic electromotility that would induce a difference from fluid motion (21, 56) (for a more detailed intra-organ model including the cells and tectorial stucture, see *Steele, 1996* (39)). An overlay of where the organ would lie is shown for visualization purposes. These images are of a zoomed-in portion of the simulated chamber near the organ; the total modeled chamber spans 500 *μm* transversely and radially, with the organ centered at the bottom.

We find that the transverse fluid velocity is asymmetric, with the medial and lateral sides of the organ moving opposite one another. This is consistent with the rocking motion seen in Figure 3 and Figure 4G, as we supposed these measurements were primarily transverse in nature.

The radial velocity, on the other hand, is uniform in direction throughout the organ. As this is the hair-bundle-stimulating direction of fluid flow, we can interpret this as meaning that all hair bundles in a given cross-section are deflected in-phase. (Whether this deflection corresponds to an increase or decrease in transepithelial current is dependent on the hair bundle orientation. In this animal, there is variability in whether bundles point medially or radially, even within a single cross-section (40, 57). That is, in-phase deflection does not necessarily mean in-phase excitation.)

The relationship between the two phenomena can be revealed by observing the modeled motion of the BM, shown at an exaggerated scale in Figure 5E. The displacement of the BM is lopsided as a result of the asymmetry in its mechanical properties. The peak of this displacement pattern lies lateral to all hair cells. Observing the normal vectors to the BM explains the medial-pointing radial fluid velocity across the organ (Fig. 5D). By conservation of mass, the large transverse fluid displacement on the lateral side is compensated by displacement in the opposite direction medial to the peak, explaining the rocking motion (Fig. 5C).

## DISCUSSION AND CONCLUSIONS

### Interpretation of results and a hypothesized mechanism

We identified a lack of tissue-scale mechanical tonotopy in the basilar papilla of the tokay gecko, as well as an antiphase movement pattern within the organ. This is consistent with both past experimental data from other reptile species (28, 32) and computational modeling efforts (34, 39). In comparison with these previous studies, we achieved a relatively comprehensive picture of these mechanics – using OCT, we showed that these results are robust across position within the organ in all three dimensions, and consistent between *ex vivo* and *in vivo* recordings.

As tuning is present in nerve fiber responses but not observable in the mechanics at this scale, it likely arises from 1) the tectorial structures – the sallets and tectorial curtain – which are not visible in our preparation (see “Open questions and limitations of the study” below), or 2) a cellular mechanism. As for the former, these structures likely impact tuning (34, 39), but as many reptiles (e.g. the alligator lizard) do not have tectorial structures (29, 30, 40), they cannot be necessary for the manifestation of tonotopy *per se*. As for cellular mechanisms, electrical tuning may take place at basal (low-frequency) regions, but are unlikely to be effective above 1 kHz in this species (4–9, 13).

We hypothesize that tuning occurs at the hair bundle level, with tissue-scale mechanics being responsible for providing fluid velocity in the proper direction to stimulate the bundles (by untuned rocking), thereby activating the tuned mechanism. For example, it has been proposed that the stereocilia and tectorial structures together form a mechanical resonator that changes in resonant frequency along the length of the cochlea, with tonotopy remaining even when the tectorial structures are not present (13, 34, 57). As hair bundles contain the motor protein myosin (58–60), they may also employ active mechanical amplification and sharpening of tuning (61–65).

### Implications for other species

The results in Figures 2 and 3 show that the untuned rocking mechanism is present at both high- and low-frequency regions, and in regions both with and without sallets. Combined with previous data from the alligator lizard (32–36), which has free-standing hair bundles rather than sallets or a tectorial curtain, this indicates a common principle at play which could generalize across a range of species. With respect to tissue-scale mechanical tuning, we note two (non-exhaustive) factors that may explain its absence in this organ, and relate them to the organs of other species:

First, the tokay gecko cochlea is short in comparison to mammalian cochleae – only *~1*.7 mm (40) *–* which precludes the presence of a cochlear traveling wave (15). Most reptiles, and even some birds known to have excellent hearing such as the budgie parrot (*Melopsittacus undulatus*), have cochleae shorter than or of similar length to that of the tokay gecko (30, 40), and thereby must also achieve frequency selectivity without a traveling wave. In owls, on the other hand, the cochlea can be as long as 11 mm, similar in length to those of gerbils (30, 66); in these longer cochleae, a traveling wave manifests (15, 67) and tissue-scale tuning is more likely to be present. Cochlear length alone is not a sufficient condition, however: *in vivo* displacement responses in the chicken basilar papilla show that while the organ sustains a traveling wave, it does not exhibit significant tissue-scale mechanical tonotopy (67).

Second, the organ has neural tuning to low frequencies at the base and to high frequencies at the apex (opposite to mammals and birds), but the BP is larger in cross-sectional area and radial width at the apex than at the base (the same as in mammals and birds) (40). This would lead one to believe that the tissue is locally stiffer in its low-frequency regions, which would be counter-productive in producing mechanical resonance consistent with neural tuning. This same stiffness gradient oddity is present in many reptile species (40).

As for the rocking motion, we explain this phenomenon with a two-beam model (Fig. 5) that depends only on an asymmetry in the BM, which appears to be present across animals in the class *Reptilia* – it appears in most lizards, snakes, sphenodon (tuatara), testudines, and very strikingly so in crocodilians (29, 30, 40). In the most extreme case, such as at some positions in the cochlea of the red-eared slider turtle (*Chrysemys scripta elegans*), the hair cells do not even lie on the basilar membrane but instead on the stiff cartilage medial to it; a cross-section of this organ is shown in Figure 5F. This can be seen as a degenerate case of our model, in which the stiffness on the medial side (previously interpreted as the fundus) is effectively infinite. The schematic below the anatomical diagram shows how hair bundle stimulation might arise in such an architecture.

It is thus reasonable to extend our hypothesis – that tuning arises at the hair bundle level, and the organ rocks in an untuned manner to activate this process – to a variety of reptilian species. In longer cochleae with directionally opposite stiffness gradients, this mechanism may be present *alongside* tissue-scale mechanical tuning. Future studies in turtle and crocodile cochleae would be particularly valuable to interrogate this.

### Open questions and limitations of the study

The majority of the presented data were collected from excised cochleae, and thus may be limited to the passive mechanics of the organ. Should tonotopy manifest in the tissue-scale mechanics of the BP *in vivo*, synthesis with the present study would imply that it is the result of an active process (likely at the hair bundle level). However, our *in vivo* findings do not indicate the presence of tonotopy, and previously measured basilar membrane responses in alligator lizard *in vivo* did not show tonotopy (26), so this is improbable.

We also cannot be sure if the tectorial structures are present in our preparations; the resolution of the device and translucency of the curtain and sallets make this impossible to assess. Even in our *in vivo* images (Fig. 4), where these structures should be intact, we could not resolve these structures. While the removal of Reissner’s membrane in our *ex vivo* experiments could result in damage to the curtain, it is unlikely that it would destroy it along the entire length of the organ. We thus presume the tectorial structures are present, but avoid making any claims regarding their motion.

A multimodal approach may be required for precisely distinguishing the responses of tectorial structures, discriminating innervated vs. non-innervated cells (24), or determining the stimulus phase relative to hair bundle polarity (40). For example, a method could be designed in which positions of OCT measurements are registered to anatomical structures *post hoc* with a higher-resolution imaging technology. Further examination of the gecko cochlea in these ways could offer new insight into the role that hair bundle mechanics plays in auditory organs, which remains one of the most elusive features of the sensation of sound.

## METHODS

### Experimental Preparation

The cochlea was isolated by adapting the method described in *Chiappe et al. (2007)* (24). In each experiment, a tokay gecko (of either sex) was decapitated after being euthanized with a lethal (300 mg/kg), intracoelomic injection of sodium pentobarbital (Euthasol®). The lower jaw was removed to expose the ventral floor of the skull. The skull was immediately placed on ice, and the columella was cut near the oval window. The otic capsule was exposed by removing the soft tissue and opened widely with scalpel and forceps. The cochlear duct, including the basilar papilla and lagena, was gently isolated after cutting away the eighth cranial nerve, semicircular canals, utricle, and saccule. The isolated cochlear duct was placed in artificial perilymph solution containing (in mM) 175 Na^+^, 3 K^+^, 2 Ca^2+^, 1 Mg^2+^, 177 Cl^-^, 1 HPO_4_^2-^, 5 HEPES, 3 pyruvate, 10 D-(+)-glucose (68–70). The other otic capsule was kept in ice and opened (within 30 minutes) in case the first cochlea failed to show vibratory responses to acoustic stimuli.

To isolate the fluid compartments of the cochlea, we mounted the cochlea on a plastic disc with an almond-shaped opening at the center. The length and width of the opening were adjusted to fit the organ so that the round window opening lied entirely on one side of the disk and the oval window opening lied on the other side. The apical and basal extremes of the organ were placed in parallel with the length of the hole. To prevent any leakage, we sealed the sides of the organ with cyanoacrylate adhesive; it was ensured that the adhesive did not enter the fluid spaces of the cochlea. Reissner’s membrane was then removed using forceps.

The disc was placed within an experimental chamber adapted from *Alonso et*.*al*. (2025) (19, 71) (see also Figure 1C). The chamber further from the OCT device adjoined the round window opening of the organ, and was in contact with the perilymph solution to simulate scala tympani. This compartment was near the speaker delivering the acoustic stimuli. The other compartment adjoined the side of the hair cells and was in contact with artificial endolymph solution to simulate scala media. The endolymph solution contained (in mM, except for Ca^2+^) 3 Na^+^, 175 K^+^, 26 *μ*M Ca2^+^, 1 Mg^2+^, 175 Cl^-^, 1 HPO_4_^2-^, 5 HEPES, 3 pyruvate, 10 D-(+)-glucose(68–70). The chamber was distinct to that of *Alonso et*.*al*. (2025) as it did not include electrodes for setting endocochlear potential or recording cochlear microphonic.

The chamber was mounted on a two-axis goniometer (SWEBO Gon, Shanghai shebao Optoelectronic Technology Co., Ltd., Shanghai, China), and then on the optical bench as described by *Alonso et*.*al. (2025)* (71).

### Stimulation

Zwuis multitone stimuli (one second in duration, comprising 13 tones between 531-6078 Hz) were created using custom-written LabVIEW (LabVIEW 2019, National Instruments, Austin, TX) and Matlab (Matlab R2021a, MathWorks, Natick, MA) programs(45, 46). After signal conditioning in a power amplifier (SA1, Tucker-Davis Technologies, Alachua, FL), acoustic stimuli were generated by an earphone (ER-3C, Etymotic Research, Etymotic Research, Elk Grove Village, IL) coupled to the innermost compartment of the experimental chamber. The sound pressure in the air cavity was measured throughout each experiment with a free-field microphone (4939-A-011, Hottinger Bruël & Kjær, Marlborough, MA), and was ensured to lie between 75 and 90 dB SPL at all frequencies and recorded positions.

Note that these stimuli are presented directly to the cochlea rather than traveling through the entire outer and middle ear pathway. Thus, an 80 dB SPL stimulus in our preparation is not equivalent to an 80 dB stimulus presented outside of an intact gecko’s head. For example, if the outer and middle ear provide 20 dB of gain, our stimulus is equivalent to if a gecko were presented a 60 dB stimulus externally.

### *In vivo* stimulation

In the case of *in vivo* OCT experiments, we placed the speaker near the tympanic membrane and applied Zwuis multitone stimuli (one-second duration, 80 dB sound comprising 13 tones between 673-6532 Hz).

### Optical Coherence Tomography (OCT)

To evaluate the condition of each preparation and measure its mechanical response to sound *ex vivo*, we used an OCT device with an 880 nm center wavelength (GAN621, Thorlabs, Newton, NJ). The imaging resolution of the device is 2.8 *μm* in the lateral direction and 4.4 *μm* in the axial direction. The resolution of displacement measurements varied by frequency (see “Analysis of Cochlear Responses” below), but was generally on the order of 0.2 nm to 1 nm. Further details of the optical setup are described by Gianoli, Alonso *et al*. (19, 71).

To measure mechanical response to sound *in vivo*, we used an OCT device with an 1300 nm center wavelength (TEL321, Thorlabs, Newton, NJ). The imaging resolution of the device is 13 *μm* in the lateral direction and 5.5 *μm* in the axial direction.

Once the cochlea was mounted (*ex vivo*) or round window was opened (*in vivo*), ThorImage (ThorLabs) was used to observe live *en face* video images and at-depth images (B-Scans, as seen in Figs 1, 2, 3 and 4) to orient ourselves within the cochlea. After assessing the condition of the preparation, cross-sections of high visibility were found and the goniometer was adjusted to obtain maximal signal-to-noise ratio in the images. The approximate percent distance from the base of these cross-sections were recorded.

Further images and displacement measurements were then acquired using LabVIEW and Matlab routines. Ten evenly-spaced axial lines along the organ were selected for displacement measurement. At each line, we recorded an OCT M-Scan – a time series of axial scans from which vibration profiles can be derived along the entire axial line – with a sample rate of 100 kHz as the stimulus was presented (see “Stimulation” above). B-Scans are median filtered and Gaussian filtered. Adobe Photoshop was used to paint fluid black in the scans to enhance readability.

### Analysis of Cochlear Responses

M-Scans, acquired as described in the preceding subsection, were processed in Matlab software to derive the frequency-domain displacement responses at each point along the ten axial scans (using the method of spectral domain phase microscopy(43)). Magnitudes and phases of these responses were observed point-by-point to generate the data shown in Figs 2, 3 and 4.

The criterion for discarding low-quality signals was determined by an amplitude-based thresholding at each point and each stimulus frequency. We compared the magnitude of the measured response at that frequency and position (signal) to the mean and standard deviation of the magnitudes of the measured responses at the same position in the ten frequency bins above and below the stimulus frequency (noise). If the signal was less than two standard deviations above the mean noise level, the value was considered insignificant.

The amplitude-phase maps of Figure 2 A, C and E are generated by isolating the measured positions within the organ by hand, determined by comparison with the B-Scan at the same position. Insignificant points were marked as having amplitude 0, and the map was Gaussian-filtered to provide smooth images. No such filtering is used for the presented data in Figures 2 and 3 B, D and E.

### Auditory Brainstem Response (ABR) Recording

Animals were anesthetized with an intracoelomic injection of ketamine (14 mg/kg; 010177, Covetrus, Portland, ME) and dexmedetomidine (0.14 mg/kg; 1457092, Henry Schein Medical, Melville, NY). If the animal became active before the recording session was completed, an additional one-third of the original anesthesia dose was injected. Once a suitable plane of anesthesia was reached (assessed via righting reflex), the anesthetized animal was placed in a sound-isolated, electrically-shielded box (72). Needle electrodes (GRD-SAF, The Electrode Store) were then subdermally placed behind the tested ear (reference electrode), in the scalp between the ears (active electrode), and in the back near the tail (ground electrode). ABRs were evoked by tone bursts of 0.5, 2.55, and 5 kHz produced by an open-field magnetic speaker connected to a power amplifier (MF1 and SA1, Tucker-Davis Technologies). Each 5 ms burst was presented 33.3 times per second with alternating polarity. The onset and offset of each burst was tapered with a squared cosine function. For each frequency, the sound pressure level was lowered from 82 dB SPL in 5–10 dB steps until the threshold was reached. If 82 dB SPL was not enough to elicit a response, higher intensities were produced. The speaker was calibrated with a ¼ inch condenser microphone (4939-A-011 and 2690-A-0S1, Bruel and Kjær) placed 5 cm away from the speaker. The electrical response evoked by the tone bursts and measured by the needle electrodes was amplified 10,000 times and bandpass filtered at 0.3–3 kHz (P55, Astro-Med Inc.). The amplified response was then digitally sampled at 10 ^µ^s intervals with a data acquisition device (PCIe-6353, National Instruments) controlled by custom software (LabVIEW 2019, National Instruments). The electrical responses to 1,000 bursts were averaged at each intensity level to determine the threshold, which was defined as the lowest level at which any response peak was distinctly and reproducibly present. Visual inspection of the vertically stacked responses facilitated threshold determination. ABR measurements were taken before and after surgery and compared to ensure hearing integrity of the organ before *in vivo* OCT recordings were taken.

### Mathematical Model

The two-beam model of the BM (Fig. 5A) is described by solving one dynamic beam equation for each beam, and boundary conditions ensuring continuity between the two beams at their point of contact. The dynamic beam equation in the phasor domain is given by

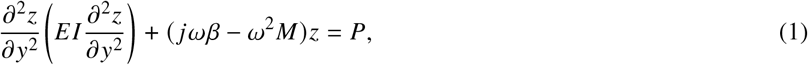

where *z* is the transverse displacement of the beam (“up-down” in Figure 5), *y* is the radial displacement across the width of the beam (“left-right”), *E* is the Young’s modulus of the beam, *I* is its moment of inertia, *β* is the resistance component of its impedance, *M* is the mass component, and *P* is the magnitude of the sinusoidal pressure stimulus with radian frequency *ω* uniformly applied on the surface of the beam.

Each of the two beams (fundus labeled *A*, flexible BM labeled *B)* has a distinct displacement profile *z*_*A,B*_, *E*_*A,B*_, *I*_*A,B*_, length *L*_*A,B*_. That is, this is a system of two fourth-order differential equations, requiring four boundary values to achieve a unique solution.

The beams are assumed to be simply supported at their junctions with the limbic cartilage (*y* = 0, *y* = *L*_*A*_ + *L*_*B*_) and continuous in value and first three derivatives with the flexible BM at its radial end *y* = *L*_*A*_:

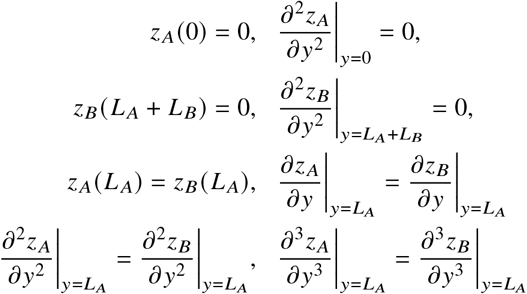

This system of equations is solved analytically by the method of undetermined coefficients; solution details and parameters are presented in the Supplementary Information.

To determine the fluid velocity patterns shown in Figure 5C-D, the Laplace equation was solved in a rectangular region (width *W*, height *H)* assuming the beam (moving as determined by solution to the system of equations above) lied at the center of the bottom boundary. We assumed the two-dimensional fluid velocity potential *ϕ* satisfies Laplace’s equation

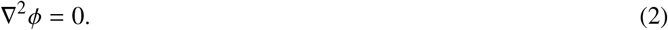

Recall that *∇ϕ* = (*v*_*y*_ *v*_*z*_)^T^, the *y* and *z* components of fluid velocity. We impose the natural boundary conditions, i.e. that fluid cannot leave the sides of the region:

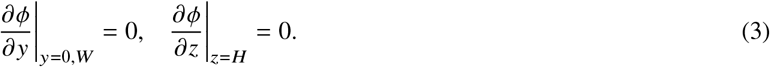

The boundary condition on the bottom surface, where the *z-*derivative of *ϕ* should be the velocity of the beam, is

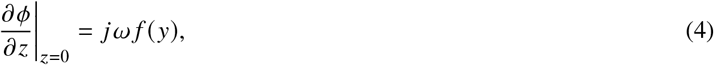

where

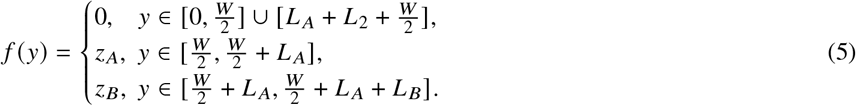

The *jω* represents the time derivative in the phasor domain.

This is a standard von Neumann problem, which can be solved by the method of separation of variables (73). The solution is

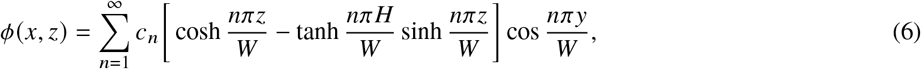

where *c*_*n*_ is a scaled cosine series coefficient for the velocity of the beam:

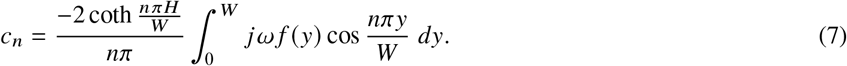

Finally, the presented velocities are found to be the components of *∇ϕ*. We use the first 100 components of the cosine series to generate the data shown in Figure 5C-D.

## DATA AVAILABILITY STATEMENT

The corresponding data sets supporting the findings in this manuscript have been deposited at Zenodo under the DOI: https://doi.org/10.5281/zenodo.22097763 and are publicly available as of the article publication date.

## AUTHOR CONTRIBUTIONS

Conceptualization, B.L.F. and A.J.H.; methodology, B.L.F., Y.V., K.H., B.A.F., A.J.H.; investigation, B.L.F., Y.V., K.H., B.A.F., A.J.H.; writing-–original draft, B.L.F.; writing-reviewing & editing, B.L.F., Y.V., K.H., B.A.F.; funding acquisition, A.J.H.; resources, B.L.F., Y.V., K.H., B.A.F., A.J.H.; supervision, A.J.H.

B.L.F., Y.V., K.H. contributed equally to this work. A.J.H. is deceased and did not read the final version of this manuscript.

## ACKNOWLEDGMENTS

We acknowledge the Comparative Bioscience Center for anesthetic assistance and support. We also acknowledge Marcel van der Heijden for his assistance in developing our OCT displacement response acquisition software.

This work was funded by the Howard Hughes Medical Institute (B.L.F., Y.V., B.A.F., A.J.H), Rockefeller University (K.H.), Naito Foundation (K.H.) and Takeda Science Foundation 2024047242 (K.H.). The authors thank all members of the Hudspeth lab, as well as Cori Bargmann and Pascal Martin, for their support and feedback on the manuscript.

We would like to dedicate this paper to the memory of A.J.H.; we are forever grateful for his mentoring and support.

## Footnotes

1 There is inconsistency in the literature on the nomenclature of reptile inner ear anatomy. We have chosen to use anatomical terms consistent with our group’s previous work on this organ (24, 25): in this work, the entire hearing organ is the *cochlea*, and the partition between scalae within the cochlea, containing hair cells, supporting cells, the basilar membrane, and tectorial structures, is the *basilar papilla*.

2 We are directly stimulating the cochlear fluid in our *ex vivo* experiments, so this is not comparable to an 80 dB SPL stimulus at the ear canal (47). The pressure gain from ear canal to fluid pressure is unknown in this species, but we can safely say the effective ear canal stimulus pressure would be lower. As we are concerned with general displacement patterns rather than specific values, this does not affect the interpretation of our results.

3 When viewing the organ with OCT *in vivo*, images appear “upside-down” relative to the previous figures due to our surgical approach. A schematic of the organ at this angle is presented in Figure 4D.

