## Supplemental Information for "Rocking Without a Tune: *Ex vivo* and *in vivo* Responses to Sound in the Basilar Papilla of the Tokay Gecko"

Brian L. Frost, Yuriria Vázquez, Kazuhiro Horii, Brian A. Fabella, and A. J. Hudspeth

### More Examples of Rocking Motion in the Basilar Papilla

In the main text we show three examples of motion maps indicating a half-cycle phase shift between abneural and neural hair cell displacement, indicative of a rocking motion. In two cases, we were able to obtain such information across a longitudinal span of the cochlea, giving the curves from main text Fig 3D and F.

More often, high visibility and thus signal-to-noise ratio is only feasible at one cross-section, and such curves cannot be derived. In this case, we can observe the motion map within each cross-section as in main text Fig 3A, C and E. We saw the half-cycle shift in at least one cross-section of $N = 23$ organs. (Other preparations did not possess sufficient visibility to measure across an entire cross-section, but still allowed for assessment of tonotopy.) In Fig S1, we show examples of this phenomenon in eight additional organs to drive home the robustness of this result. Note that positions from the apex, center, and base of the organ are all represented.

### More Examples of *in vivo* Responses Showing Lack of Tonotopy

In the main text we show one example of *in vivo* displacement responses from TK 99, which showing both a lack of tonotopy (Figure 4F) and rocking motion (Figure 4G) consistent with our *ex vivo* results. We were able to achieve statistically significant magnitude responses in three other animals at the neural side of the BP. These results are shown in Figure S2. The responses are low-pass, similar to those from the *in vivo* (Figure 4) and *ex vivo* (Figure 2) responses.

### Two-Beam Model Formulation and Solution

The two-beam model of the BM (main text Figure 5) is described by one dynamic beam equation for each beam, and boundary conditions ensuring continuity between the two beams at their point of contact. The dynamic beam equation in the phasor domain is given by

$$\frac{\partial^2 z}{\partial y^2}\left(EI\frac{\partial^2 z}{\partial y^2}\right) + (j\omega\beta - \omega^2 M)z = P, \tag{S1}$$

where $z$ is the displacement of the beam ("up-down" in Fig 4E), $y$ is the displacement across the width of the beam ("left-right" in Fig 4E), $E$ is the Young's modulus of the beam, $I$ is its moment of inertia, $\beta$ is the resistance component of its impedance, $M$ is the mass component, and $P$ is the

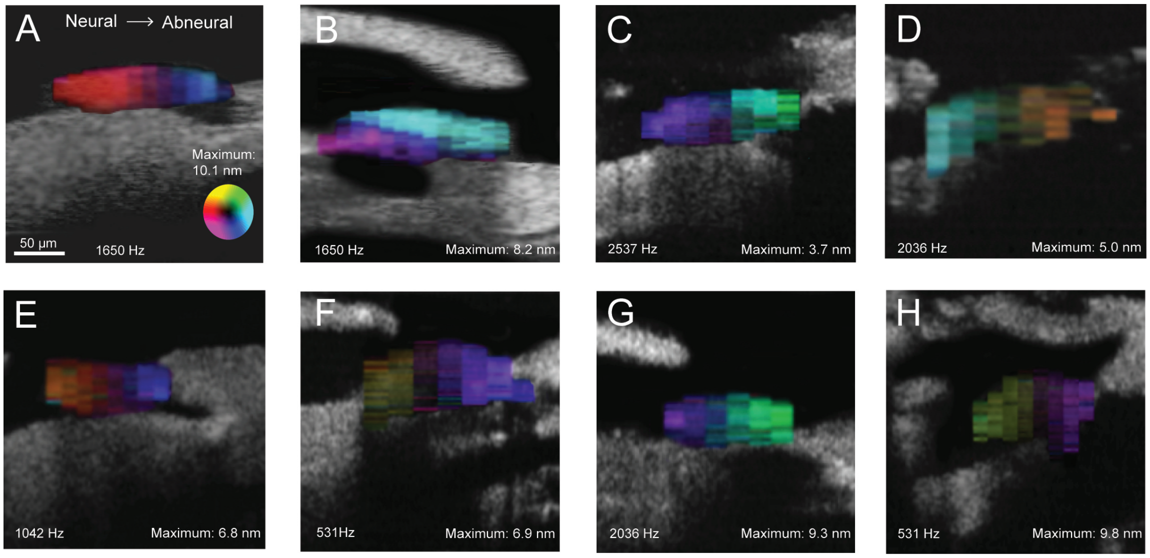


Figure S1: Motion maps of the BP in eight different animals showing the characteristic rocking motion seen in the data of main text's Figure 3 and the model results of Fig 4G; maps are read identically to those of main text's Figure 3. Stimulus frequencies are given within each panel. **A** – TK 17, basal; **B** – TK 18, basal; **C** – TK 88, central; **D** – TK 85, apical; **E** – TK 92, basal; **F** – TK 90, apical; **G** – TK 74, central; **H** – TK 77, apical. The convex bend on the top of panel H is a floating piece of Reissner's membrane, which has been deliberately torn.

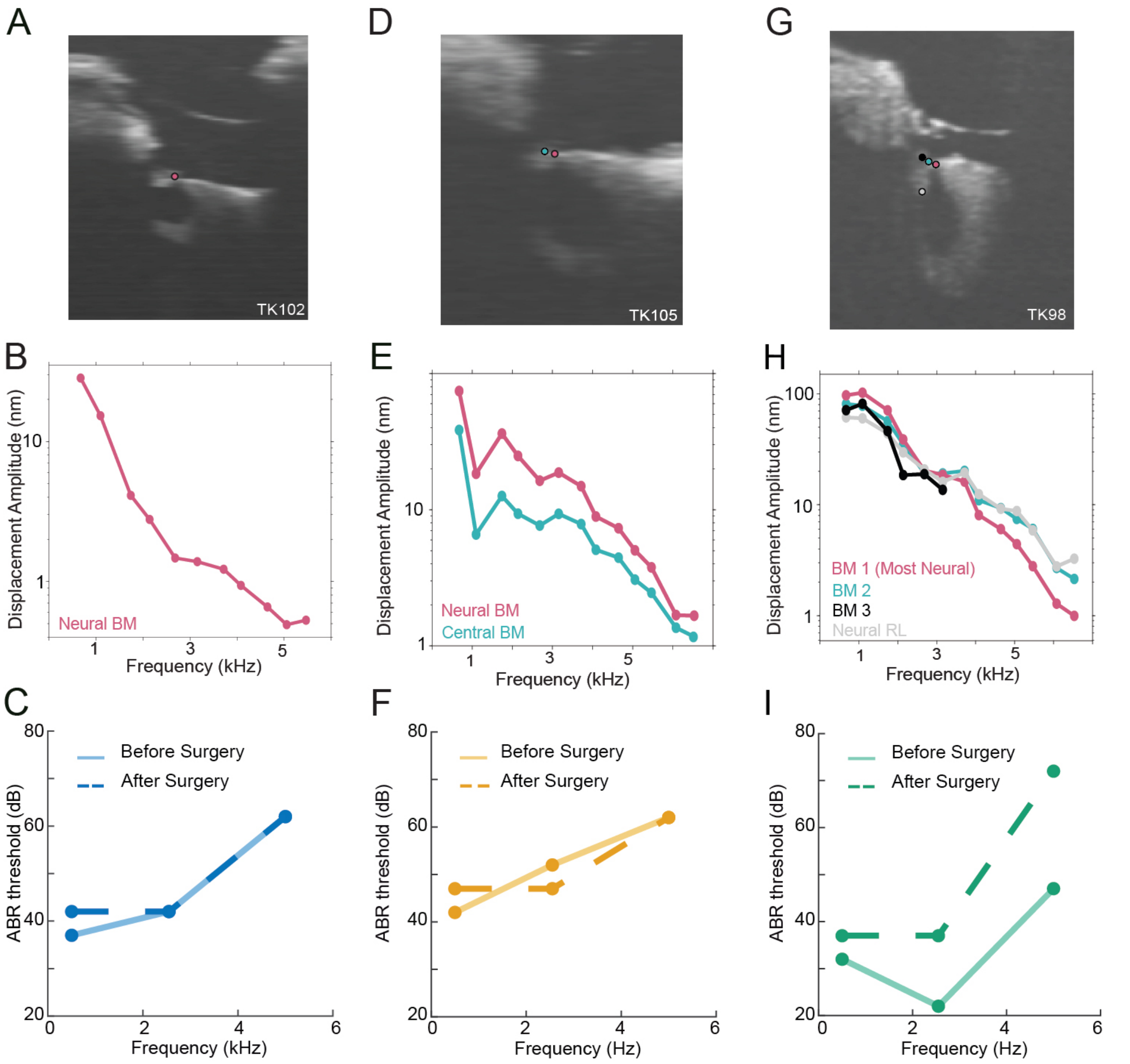


Figure S2: Displacement amplitude responses in three tokay gecko cochleae *in vivo* showing low-pass responses, consistent with main text's Figure 4F. **A** – OCT image from TK 102, with a colored circle marking a neural BM position from which data was collected; **B** – Magnitude response at the position marked in A; **C** – ABR thresholds before and after surgery for Tk102. **D-F** Same as A-C but in TK 105, with measurements at two neural BM positions; **G-I** – Same as A-B but in TK 98, with measurements at two neural and one central BM positions and one neural reticular lamina position.

magnitude of the sinusoidal pressure stimulus with radian frequency $\omega$ uniformly applied on the surface of the beam.

Each of the two beams (fundus labeled $A$, flexible BM labeled $B$) has a distinct displacement profile $z_{A,B}$, $E_{A,B}$, $I_{A,B}$, length $L_{A,B}$. That is, this is a system of two fourth-order differential equations, requiring four boundary values to achieve a unique solution.

The beams are assumed to be simply supported at their junctions with the limbic cartilage ($y = 0$, $y = L_A + L_B$) and continuous in value and first three derivatives with the flexible BM at its radial end $y = L_A$:

$$\begin{aligned} z_A(0) = 0, &\quad \left.\frac{\partial^2 z_A}{\partial y^2}\right|_{y=0} = 0, \\ z_B(L_A + L_B) = 0, &\quad \left.\frac{\partial^2 z_B}{\partial y^2}\right|_{y=L_A+L_B} = 0, \\ z_A(L_A) = z_B(L_A), &\quad \left.\frac{\partial z_A}{\partial y}\right|_{y=L_A} = \left.\frac{\partial z_B}{\partial y}\right|_{y=L_A} \\ \left.\frac{\partial^2 z_A}{\partial y^2}\right|_{y=L_A} = \left.\frac{\partial^2 z_B}{\partial y^2}\right|_{y=L_A}, &\quad \left.\frac{\partial^3 z_A}{\partial y^3}\right|_{y=L_A} = \left.\frac{\partial^3 z_B}{\partial y^3}\right|_{y=L_A} \end{aligned}$$

### Solution

The method of undetermined coefficients is used to find a particular solution for each of $z_A$ and $z_B$. As the nonhomogeneity of Eqn S1 is a constant $P$, a reasonable ansatz for a particular solution would be a constant. Assuming $\beta$ and $M$ are the same for both beams gives the same particular solution for $z_1$ and $z_2$:

$$z_p = \frac{P}{j\omega\beta - \omega^2 M}. \tag{S2}$$

Assuming $E$ and $I$ are constant, these are fourth-order linear differential equations with constant coefficients; we use the method of the auxiliary equation to solve for the associated homogeneous solutions. In the case of $z_A$, the auxiliary polynomial is

$$n^4 = -\frac{j\omega\beta - \omega^2 M}{E_A I_A}. \tag{S3}$$

Thus, the associated homogeneous solution is a linear combination of exponentials where the exponential constants are the four complex fourth-roots of the above ratio, and the total solution is the sum of this and the particular solution:

$$z_A = \frac{P}{j\omega\beta - \omega^2 M} + \sum_{i=1}^{4} C_i e^{n_i y}, \quad C_i \in \mathbb{C}, \quad n_i = \left(\frac{-(j\omega\beta - \omega^2 M)}{E_A I_A}\right)^{1/4}. \tag{S4}$$

The solution for $z_B$ takes the same form, but the constants differ as $E$ and $I$ differ:

$$z_B = \frac{P}{j\omega\beta - \omega^2 M} + \sum_{i=1}^{4} D_i e^{m_i y}, \quad D_i \in \mathbb{C}, \quad m_i = \left(\frac{-(j\omega\beta - \omega^2 M)}{E_B I_B}\right)^{1/4}. \tag{S5}$$

Finally, we can solve for the unknown coefficients $C_i$ and $D_i$ simultaneously by application of the boundary conditions above (shifting $y$ coordinates for $z_2$ by $L_A$ to simplify notation). The resulting $8 \times 8$ system to be solved is:

$$\begin{pmatrix} 1 & 1 & 1 & 1 & 0 & 0 & 0 & 0 \\ n_1^2 & n_2^2 & n_3^2 & n_4^2 & 0 & 0 & 0 & 0 \\ 0 & 0 & 0 & 0 & e^{m_1 L_B} & e^{m_2 L_B} & e^{m_3 L_B} & e^{m_4 L_B} \\ 0 & 0 & 0 & 0 & m_1^2 e^{m_1 L_B} & m_2^2 e^{m_2 L_B} & m_3^2 e^{m_3 L_B} & m_4^2 e^{m_4 L_B} \\ e^{n_1 L_A} & e^{n_2 L_A} & e^{n_3 L_A} & e^{n_4 L_A} & -1 & -1 & -1 & -1 \\ n_1 e^{n_1 L_A} & n_2 e^{n_2 L_A} & n_3 e^{n_3 L_A} & n_4 e^{n_4 L_A} & -m_1 & -m_2 & -m_3 & -m_4 \\ n_1^2 e^{n_1 L_A} & n_2^2 e^{n_2 L_A} & n_3^2 e^{n_3 L_A} & n_4^2 e^{n_4 L_A} & -m_1^2 & -m_2^2 & -m_3^2 & -m_4^2 \\ n_1^3 e^{n_1 L_A} & n_3^2 e^{n_2 L_A} & n_3^3 e^{n_3 L_A} & n_4^3 e^{n_4 L_A} & -m_1^3 & -m_2^3 & -m_3^3 & -m_4^3 \end{pmatrix} \begin{pmatrix} C_1 \\ C_2 \\ C_3 \\ C_4 \\ D_1 \\ D_2 \\ D_3 \\ D_4 \end{pmatrix} = \begin{pmatrix} -z_p \\ 0 \\ -z_p \\ 0 \\ 0 \\ 0 \\ 0 \\ 0 \end{pmatrix}$$

The constants are solved for by matrix inversion; $z_A$ and $z_B$ are thus determined.

### Parameter Selection

Parameter values used for the model are given in Table S1. The geometric parameters are derived from *Wever (1978)*[1], and the mass and damping parameters $(M, \beta)$ are presumed to be similar to the human[2].

As for Young's modulus, we choose that of the fundus to be higher than the human BM's, and then assume the modulus of the flexible BM is half that much. These values are found by fitting, and we see that it is the ratio $E_A/E_B$ that has the most effect on the main result (this is intuitive, as the ratio is a quantification of BM asymmetry). Further experiments would be necessary to determine these values more precisely.

Geometric parameters are also used to compute the moment of inertia, $I$, according to the standard formula:

$$I_{A,B} = \frac{L_{A,B} h_{A,B}^3}{12}, \tag{S6}$$

where $h$ is the thickness of the beam.

### Fluid Velocity

To produce the main text's Fig 4G, we solved for the fluid velocity in a rectangular region (width $W$, height $H$) assuming the beam, moving as solved for above, lied at the center of one boundary. We assumed the two-dimensional fluid velocity potential $\phi$ satisfies Laplace's equation

$$\nabla^2 \phi = 0. \tag{S7}$$

| Parameter | Fundus | Flexible BM |
|---|---|---|
| $L_{A,B}$ (length, $\mu$m) | 60 | 10 |
| $M$ (mass component of impedance, kg/mm) | $1.5 \times 10^{-6}$ | $1.5 \times 10^{-6}$ |
| $\beta$ (damping component of impedance, Ns/mm$^2$) | $2 \times 10^{-6}$ | $2 \times 10^{-6}$ |
| $E_{A,B}$ (Young's modulus, N/mm$^2$) | 16 | 8 |
| $h_{A,B}$ (thickness, $\mu$m) | 30 | 3 |
| $\omega$ (stimulus frequency, rad/s) | $2\pi\times$ 1000 | $2\pi\times$ 1000 |

Table S1: Parameters used to produce the model results shown in main text Fig 4G. Parameters are from *Wever (1978)*[1] and the authors' estimates inspired by *Steele and Taber* (1979)[2].

Recall that $\nabla\phi = (v_y\ v_z)^{\mathrm{T}}$, the $y$ and $z$ components of velocity. We impose the natural boundary conditions, i.e. that fluid cannot leave the sides of the region:

$$\left.\frac{\partial\phi}{\partial y}\right|_{y=0,W} = 0, \quad \left.\frac{\partial\phi}{\partial z}\right|_{z=H} = 0. \tag{S8}$$

The boundary condition on the bottom surface, where the $z$-derivative of $\phi$ should be the velocity of the beam, is

$$\left.\frac{\partial\phi}{\partial z}\right|_{z=0} = j\omega f(y), \tag{S9}$$

where

$$f(y) = \begin{cases} 0, & y \in [0, \frac{W}{2}] \cup [L_A + L_2 + \frac{W}{2}], \\ z_A, & y \in [\frac{W}{2}, \frac{W}{2} + L_A], \\ z_B, & y \in [\frac{W}{2} + L_A, \frac{W}{2} + L_A + L_B]. \end{cases} \tag{S10}$$

The $j\omega$ represents the time derivative in the phasor domain.

This is a standard von Neumann problem, which can be solved by the method of separation of variables. The full steps of the solution are omitted, as they can be found in any text on differential equations. The solution is

$$\phi(x,z) = \sum_{n=1}^{\infty} c_n \left[\cosh\frac{n\pi z}{W} - \tanh\frac{n\pi H}{W}\sinh\frac{n\pi z}{W}\right]\cos\frac{n\pi y}{W}, \tag{S11}$$

where $c_n$ is a scaled cosine series coefficient for the velocity of the beam:

$$c_n = \frac{-2\coth\frac{n\pi H}{W}}{n\pi}\int_0^W j\omega f(y)\cos\frac{n\pi y}{W}\,dy. \tag{S12}$$

Finally, the presented $y$-direction velocity is found by the relation

$$\frac{\partial\phi}{\partial y} = v_y.$$
